# Commensal bacterial-derived retinoic acid primes host defense to intestinal infection

**DOI:** 10.1101/2021.01.27.428280

**Authors:** Vivienne Woo, Emily M. Eshleman, Jordan Whitt, Seika Hashimoto-Hill, Shu-en Wu, Laura Engleman, Taylor Rice, Rebekah Karns, Bruce A. Vallance, Theresa Alenghat

## Abstract

Interactions between the microbiota and mammalian host are essential for effective defense against pathogenic infection; however, the microbial-derived cues that mediate this beneficial relationship remain unclear. Here, we find that the intestinal epithelial cell (IEC)-associated commensal bacteria, Segmented Filamentous Bacteria (SFB), promotes early protection against the bacterial pathogen, *Citrobacter rodentium*, independently of CD4^+^ T cells. Global analyses demonstrated that SFB induced histone modifications in IECs at sites enriched for the retinoic acid receptor (RAR) motif. Interestingly, SFB-colonized mice exhibited greater expression of RAR targets during infection relative to germ-free mice, suggesting SFB may enhance defense through retinoic acid (RA) signaling. Consistent with this, supplementing germ-free mice with RA decreased pathogen levels. Further, mice with impaired RA-responsiveness in IECs displayed increased susceptibility to *C. rodentium* infection. RA was elevated in the intestine of mice colonized with SFB, indicating that the presence of commensal bacteria can modulate intestinal RA levels. However, this regulation by SFB was not dependent on mammalian RA production. Sequence analyses suggested that RA-generating enzymes are expressed by a subset of commensal bacteria. Remarkably, RA was produced by intestinal bacteria including SFB, and inhibiting RA signaling blocked SFB-induced protection against *C. rodentium* infection. These data collectively reveal RA as an unexpected microbiota-derived metabolite that primes innate intestinal defense and suggests that pre- and probiotic approaches to elevate RA could prevent or combat pathogenic infection.

## Introduction

The mammalian intestine is inhabited by trillions of commensal microbes, collectively referred to as the microbiota. In addition to innocuous commensals, the gastrointestinal tract is constantly at risk of invasion and infection by pathogenic microbes. Interactions between the intestinal microbiota and the mammalian host are essential for effective defense against pathogens, as loss of the microbiota in germ-free and antibiotic-treated animals leads to increased susceptibility to enteric and non-enteric infection (Abt and Pamer, 2014; Benson et al., 2009; Ganal et al., 2012; Ivanov et al., 2009). Intestinal epithelial cells (IECs) reside at the direct interface between the host and commensal microbes and, therefore, carry the potential to critically respond to signals from the microbiota. Besides functioning as a physical barrier, these cells actively respond to microbial challenges by secreting antimicrobial peptides, mucins, chemokines and cytokines that prime and regulate innate and adaptive immunity (Gallo and Hooper, 2012; Peterson and Artis, 2014; Ramanan and Cadwell, 2016). IECs are also equipped to sense microbial stimuli through various membrane and cytoplasmic pattern-recognition receptors (Price et al., 2018).

In addition to canonical microbial sensing pathways, epigenetic mechanisms enable environmental signals to instruct cellular responses and represent another interface by which microbiota can impact mammalian cells (Amatullah and Jeffrey, 2020; Woo and Alenghat, 2017). Consistent with this, epigenetic-modifying enzymes in IECs integrate microbiota-derived signals to regulate intestinal homeostasis and immunity (Ansari et al., 2020; Navabi et al., 2017; Takahashi et al., 2009; Wu et al., 2020). Epigenetic-modifying enzymes mediate covalent chromatin modifications that alter DNA accessibility and gene expression. Thus, epigenetic modifications that are sensitive to the microbiota may identify regulatory pathways that can enhance host defense to infection (Arrowsmith et al., 2012; Kelly et al., 2018).

Increasing evidence highlights that microbiota-derived metabolites mediate the host-microbiota relationship (Lavelle and Sokol, 2020; McCarville et al., 2020; Rooks and Garrett, 2016). Commensal bacteria generate a variety of metabolites through either direct synthesis or breakdown of dietary components that can be absorbed in the intestine and potentially travel systemically (Matsumoto et al., 2018; Wikoff et al., 2009). For example, well-characterized bacterial-derived short-chain fatty acids that are produced by bacteria in the intestine can regulate local cells as well as distant tissues (Chang et al., 2014; Dalile et al., 2019; Fellows et al., 2018; Furusawa et al., 2013; Kaiko et al., 2016; Yang et al., 2020). However, despite the appreciation that commensal bacteria prime enhanced innate defenses, the underlying pathways and microbiota-derived cues that decrease host susceptibility to pathogenic infection remain poorly defined.

*Citrobacter rodentium* is a well-characterized murine bacterial pathogen that causes similar pathology to human enteropathogenic *Escherichia coli* (Mundy et al., 2005). *C. rodentium* initiates intestinal infection by adhering to the apical surface of IECs in the large intestine. Defense against *C. rodentium* requires signals from commensal microbes, as microbiota-depleted animals exhibit higher *C. rodentium* levels and impaired ability to clear the infection compared to microbiota-replete counterparts (Kamada et al., 2012; Osbelt et al., 2020; Woo et al., 2019). Segmented Filamentous Bacteria (SFB) is a commensal bacteria (Jonsson et al., 2020) that protects against a enteric pathogens such as *C. rodentium* (Chung et al., 2012; Garland et al., 1982; Heczko et al., 2000; Ivanov et al., 2009; Shi et al., 2019). Unlike the majority of commensal bacteria that are spatially separated from the epithelium, SFB directly binds to IECs in the distal small intestine (Atarashi et al., 2015; Ivanov et al., 2009; Ladinsky et al., 2019). SFB protects against *C. rodentium* infection, despite colonizing a distinct anatomical region of the intestine. Therefore, SFB likely modulates mammalian pathways rather than directly competing with *C. rodentium*, as has been shown for commensal *E. coli* and *Bacteroides thetaiotaomicron* (Kamada et al., 2012).

Previous studies have described that decreased *C. rodentium* infection in mice colonized with SFB associated with SFB-induced expansion of CD4^+^ Th17 cells that produce IL-17 and IL-22 (Goto et al., 2014; Ivanov et al., 2009). Here, we discovered that SFB also decreases initial susceptibility to *C. rodentium* infection prior to regulation by CD4^+^ T cells. ChIP-seq analyses in uninfected mice revealed that SFB colonization induced epigenetic modifications in IECs at retinoic acid receptor (RAR) motifs. Consistent with enhanced transcriptional potential, IECs from SFB-colonized mice exhibited greater induction of RAR targets relative to IECs from germ-free mice post-*C. rodentium* infection, suggesting that SFB may enhance innate defense through the RAR ligand, retinoic acid (RA). Interestingly, intestinal RA levels were increased in mice colonized with SFB and inhibiting RA signaling in SFB-colonized mice increased pathogen burden. However, SFB-dependent RA accumulation was not dependent on mammalian RA production. Instead, SFB and other commensal bacteria expressed dehydrogenase genes homologous to a microbial enzyme that converts vitamin A to RA. Remarkably, these same bacteria increased RA *in vitro* and in the intestine, thus highlighting that commensal bacteria can provide a direct source of RA that enhances defense against enteric pathogens.

## Results

### Commensal SFB protects against early infection independently of CD4^+^ T cells

*C. rodentium* is an enteric pathogen that exerts a similar pathogenesis to human enteropathogenic *E. coli* and establishes initial colonization within 2-3 days, reaching peak of infection around days 8-10 post-infection (Symonds et al., 2009). The presence of SFB in the intestinal microbiota protects against *C. rodentium* infection (Ivanov et al., 2009). Furthermore, we found that colonizing germ-free (GF) mice with only SFB was sufficient to significantly lower pathogen burdens compared to GF mice (**Figure 1A**). Interestingly, *C. rodentium* protection in SFB-colonized mice was already evident within the early phase of infection (days 3-6), suggesting that SFB may promote innate responses that decrease *C. rodentium* burden. SFB-colonization has previously been shown to induce CD4^+^ Th17 differentiation. In mice, Th17 cells activated during the peak of *C. rodentium* infection (day 10-14) mediate clearance of the pathogen by producing IL-22 and IL-17 (Ivanov et al., 2009; Omenetti et al., 2019). However, it is unclear whether Th17 cells are involved in SFB-dependent defense against *C. rodentium* during initial stages of infection. To test the involvement of Th17 cells and other CD4^+^ T helper cell populations in SFB-dependent protection against early phase *C. rodentium* infection, anti-CD4 antibodies were administered prior and during infection (**Figure 1B**) which effectively depleted CD4^+^ T cells systemically (**Figure S1A**) as well as in the colon where *C. rodentium* infects (**Figures 1C and S1B)**. Interestingly, SFB colonization led to decreased *C. rodentium* burden even when mice lacked CD4^+^ T cells (**Figure 1D)**, indicating that CD4^+^ T cells are not required for initial SFB-dependent protection.

**Figure 1:**
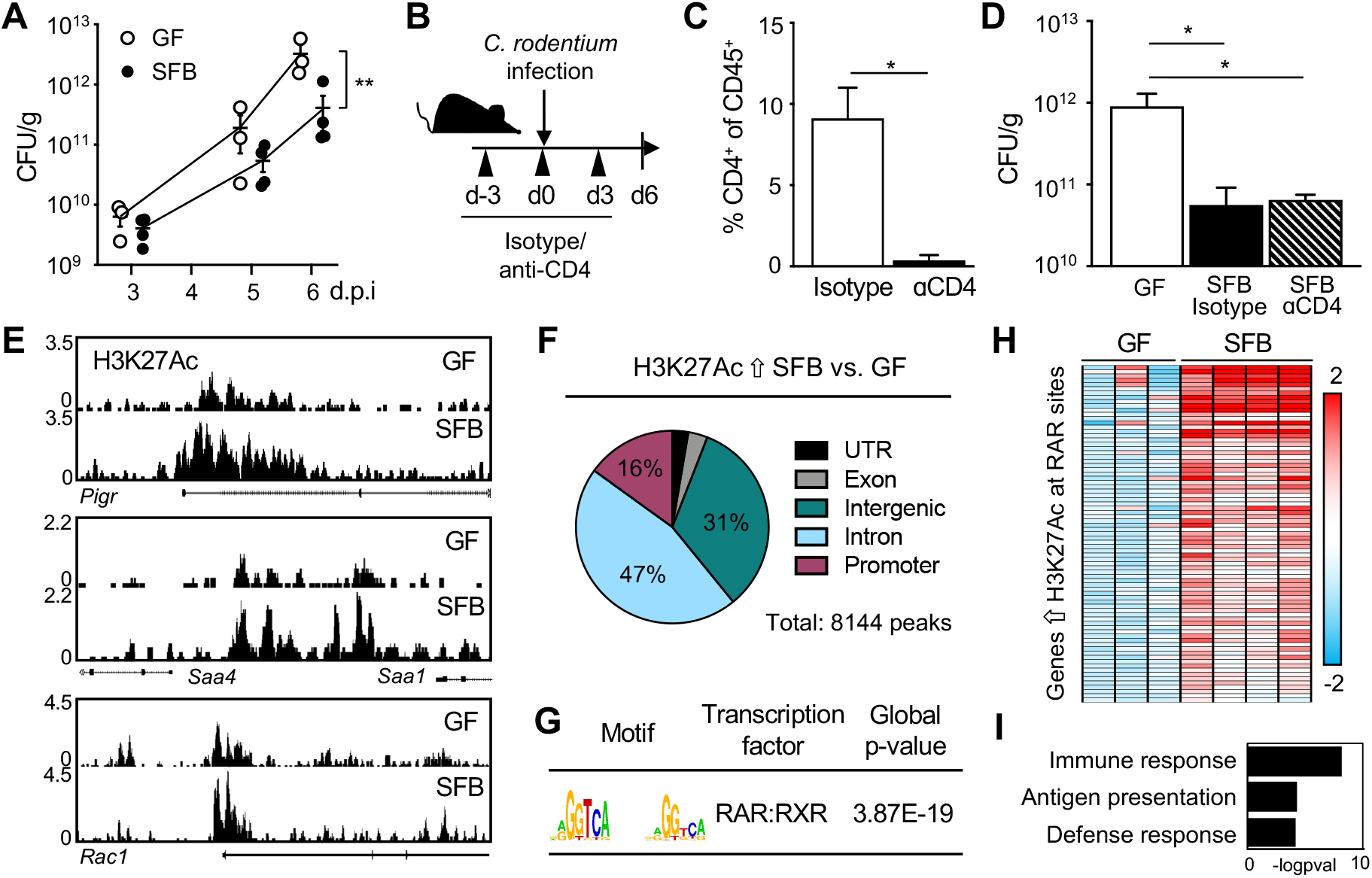
Commensal SFB primes the intestinal epithelium at retinoic acid receptor sites. (**A**) Colony-forming units (CFUs) of *C. rodentium* in stool of infected germ-free (GF) and SFB-monoassociated mice, normalized to sample weight, days 3-6 post-infection (p.i.). (**B**) Experimental approach. (**C**) Percent CD4^+^ T cells in colon from isotype and anti-CD4 treated mice. Gated on CD45^+^ cells. (**D**) *C. rodentium* CFUs in stool, normalized to sample weight, day 6 p.i. (**E**) Representative sequence tracks for H3K27Ac ChIP-seq from IECs isolated from ileum of GF and SFB-monoassociated mice, normalized to reads per million mapped reads. (**F**) Genomic distribution of H3K27Ac peaks increased in IECs of SFB versus GF mice, shown as percent of total number of differential peaks. (**G**) Motif enrichment of retinoic acid receptor (RAR) binding elements at SFB-induced H3K27Ac sites using JASPAR. (**H**) Heatmap of relative mRNA expression in IECs harvested from *C. rodentium*-infected GF and SFB mice at day 6 p.i., represented as relative fold change. (**I**) Gene ontology for RAR targets that are differentially induced in SFB-infected vs GF-infected from (G). Results are ± SEM. \**p* < 0.05, *\*\*p* < 0.01.

### Intestinal epithelium is transcriptionally primed by SFB at retinoic acid receptor motifs

Given that enhanced initial defense against *C. rodentium* was not reliant on SFB-induced CD4^+^ T cells (**Figure 1D**), we hypothesized that IECs may be important mediators of SFB-triggered defense. IECs are critically poised to respond to the microbiota and pathogens and thus play an important role in innate defense. To investigate whether SFB alters the transcriptional state of IECs, we first performed chromatin-immunoprecipitation sequencing (ChIP-seq) on primary IECs isolated from ileum of GF and SFB-monoassociated mice for the histone modification H3 lysine 27 acetylation (H3K27Ac) that characterizes primed and active chromatin (Creyghton et al., 2010). These global analyses revealed many genes that exhibit increased levels of histone H3K27Ac in response to SFB colonization, as indicated by differential peaks at multiple representative genes (**Figure 1E**). The majority of the sites with differential H3K27Ac in IECs from SFB-colonized mice occurred in regulatory gene elements (**Figure 1F)**, consistent with the known link between H3K27Ac and transcriptionally primed genes (Creyghton et al., 2010; Rada-Iglesias et al., 2011). To determine whether regions of increased histone acetylation were regulated by a shared transcription factor and/or pathway, motif enrichment analyses were performed. Sites with elevated H3K27Ac in IECs of SFB-colonized mice were significantly enriched for retinoic acid receptor (RAR) motifs (**Figure 1G**). RARs are a family of nuclear hormone receptors that regulate chromatin accessibility and gene expression by recruiting epigenetic modifiers and cofactors. Interestingly, expression of a large proportion of RAR targets with increased H3K27Ac were significantly upregulated in IECs from SFB mice during *C. rodentium* infection, compared to IECs from infected GF mice (**Figure 1H)**. Pathway analyses further revealed that the majority of the SFB-sensitive epigenetically primed RAR targets were enriched in host defense pathways (**Figure 1I**). These data demonstrate that commensal SFB modifies the epigenetic and transcriptional state of the intestinal epithelium and suggest that SFB-dependent regulation of RAR in IECs may prime innate defense against infection.

### IEC-intrinsic RAR activation enhances defense against *C. rodentium*

RARs are a family of transcription factors that bind as heterodimers with retinoid X receptors to retinoic acid response elements in the DNA. These receptors are activated by binding to the vitamin A metabolite, retinoic acid (RA). Ligand binding results in recruitment of molecular machinery that modifies local chromatin and promotes active transcription. RA and vitamin A availability can modulate *C. rodentium* and *E.coli* infection in mice and humans, respectively (Cabrera et al., 2014; McDaniel et al., 2015), provoking the hypothesis that RA may mediate the SFB-induced decrease in *C. rodentium*. Therefore, to first test whether RA is protective against *C. rodentium* in an SFB-deficient context, GF mice were treated with exogenous RA prior to and throughout the duration of infection. Administration of RA to GF mice was sufficient to protect GF mice against *C. rodentium* (**Figure 2A**), similar to protective effects described for RA-treated microbiota-sufficient mice (Snyder et al., 2019). *C. rodentium* growth and viability was not directly impaired by RA *in vitro*, indicating that decreased *C. rodentium* infection from RA supplementation in mice required the mammalian host (**Figure 2B**).

**Figure 2:**
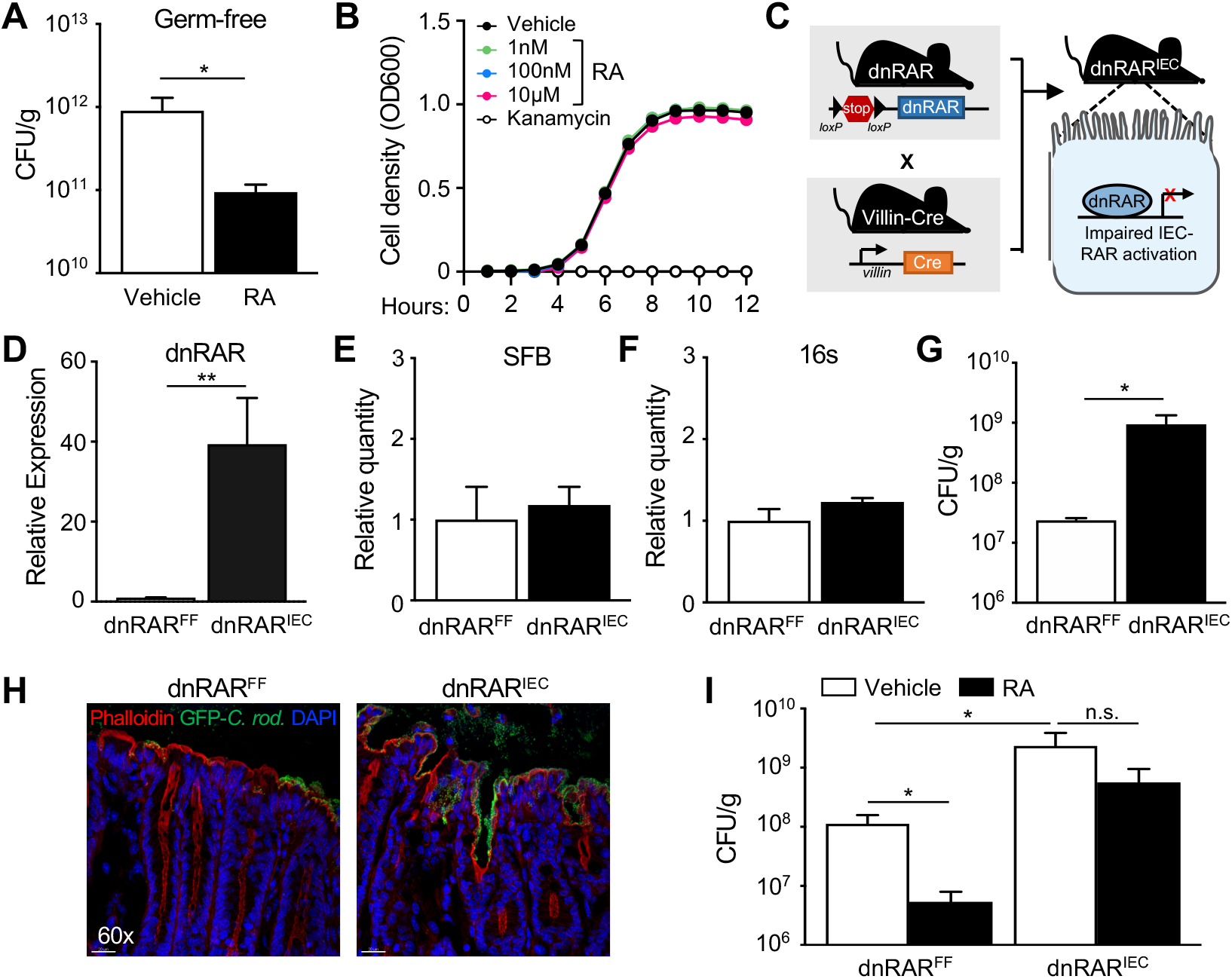
IEC-intrinsic RAR activation enhances defense against *C. rodentium*. (**A**) *C. rodentium* CFUs in stool of GF mice treated with all-trans retinoic acid (RA), day 6 p.i. (**B**) Bacterial cell density measured as the optical density at 600 nm of *C. rodentium* during growth in indicated conditions. (**C**) Control mice with a floxed stop-codon upstream of a dominant-negative RAR (dnRAR^FF^) were crossed to mice expressing IEC specific villin-cre recombinase to generate IEC-specific dnRAR expressing mice (dnRAR^IEC^). (**D**) dnRAR mRNA expression in IECs of dnRAR^FF^ and dnRAR^IEC^ mice, normalized to dnRAR^FF^. (**E, F**) Relative level of (E) SFB and (F) 16S rRNA DNA by qPCR in feces from dnRAR^FF^ and dnRAR^IEC^ mice, normalized to dnRAR^FF^. (**G**) *C. rodentium* CFUs in stool, normalized to sample weight, day 6 p.i. (**H**) Immunofluorescent staining of distal large intestine from dnRAR^FF^ and dnRAR^IEC^ mice infected with GFP-expressing *C. rodentium* (Green: GFP-*C. rodentium*, red: Phalloidin, blue: DAPI). (**I**) *C. rodentium* CFUs in stool, normalized to sample weight, day 6 p.i. Results are ± SEM. \**p* < 0.05, \*\**p* < 0.01. n.s., not significant.

The role of RA in infection has been most extensively investigated in immune cells (Hall et al., 2011a) and recent studies demonstrated that loss of RAR expression or impaired RAR responsiveness in IECs alters intestinal development and defense (Gattu et al., 2019; Iyer et al., 2020; Jijon et al., 2018). To test whether IEC-intrinsic RA signaling contributes to RA mediated protection against *C. rodentium* infection, we generated an IEC-specific dominant-negative RAR (dnRAR) transgenic mouse (**Figure 2C**) that expresses non-responsive RAR specifically in IECs (dnRAR^IEC^) (**Figure 2D**). SFB and 16S bacterial levels were similar in dnRAR^FF^ and dnRAR^IEC^ mice (**Figures 2E and 2F**). Interestingly, dnRAR^IEC^ mice exhibited significantly higher *C. rodentium* infection compared with floxed controls (dnRAR^FF^) (**Figures 2G and 2H**), a similar effect previously described for *Salmonella typhimurium* infection (Gattu et al., 2019; Iyer et al., 2020). To next decipher whether RA-driven protection requires epithelial RAR activation, control and dnRAR^IEC^ mice were treated with RA and infected with *C. rodentium*. As described above, RA-treatment decreased infection in control dnRAR^FF^ mice (**Figure 2I**); however, this RA-induced protection was greatly reduced in dnRAR^IEC^ mice (**Figure 2I**), indicating that epithelial-intrinsic RAR activity significantly contributes to RA-dependent defense against *C. rodentium*.

### SFB increases intestinal retinoic acid levels independently of host production

Our initial findings demonstrated that SFB increased histone acetylation at RAR target genes in IECs (**Figures 1E-1G**) and that the timing and magnitude of RA-induced protection against *C. rodentium* parallels the phenotype observed with SFB colonization (**Figures 1A and 2A**). Thus, to test whether SFB regulates RA in the intestine, intestinal levels of RA were compared between GF and SFB-colonized mice. Interestingly, SFB mice exhibited significantly elevated RA in intestinal contents and IECs compared to GF mice (**Figures 3A and 3B**). Consistent with increased local RA, expression of RA-sensitive genes was increased in IECs from SFB-colonized mice (**Figure 3C**). RA is generated from the vitamin A-derivative retinol in a two-step oxidation reaction involving retinol dehydrogenases (ADH, RDH) and retinaldehyde dehydrogenases (ALDH, RALDH) (**Figure 3D**). IECs are equipped to generate RA, and it was recently found that microbiota modulates intestinal tissue RA levels by regulating expression of Rdh7 in IECs (Grizotte-Lake et al., 2018). However, we found that expression of key enzymes involved in the conversion of vitamin A to RA (Aldh1a2 and Rdh7) were similarly expressed in IECs of GF and SFB mice (**Figure 3E**), suggesting that host-intrinsic enzymes required for RA production are unaffected by SFB. To further investigate the contribution of host RA synthesis on intestinal RA levels in SFB colonized mice, mice were treated with an Aldh1a2 inhibitor, WIN18446, that blocks mammalian RA production (Arnold et al., 2015). As predicted, administration of WIN18446 decreased IEC expression of mammalian RA-sensitive genes (**Figure 3F**). However, despite no effect on SFB itself (**Figure 3G**), RA levels in SFB colonized mice remained elevated with WIN18446 treatment (**Figure 3H**), demonstrating that the SFB-dependent increase in intestinal RA occurs independently of host RA synthesis.

**Figure 3:**
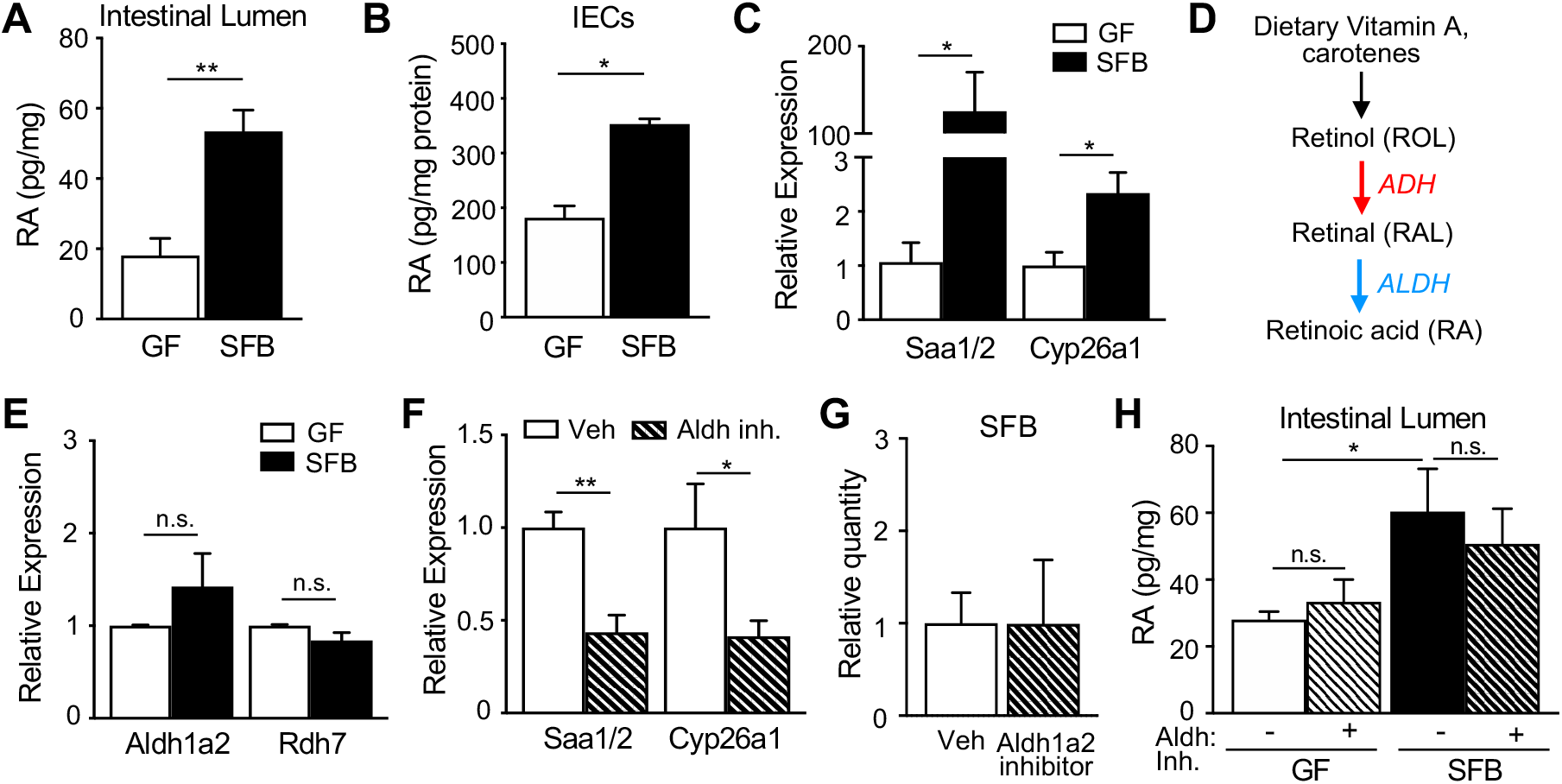
SFB increases intestinal retinoic acid levels independently of host production. (**A, B**) Retinoic acid concentration in (A) fecal samples and (B) IECs collected from GF and SFB mice, normalized to sample weight. (**C**) mRNA expression in IECs of GF and SFB mice, normalized to GF. (**D**) Schematic of mammalian vitamin A metabolism to retinoic acid. (**E**) mRNA expression in IECs of GF and SFB mice, normalized to GF. (**F**) mRNA expression in IECs of Aldh1a2 inhibitor (WIN18446) treated mice, normalized to vehicle. (**G**) Relative level of SFB DNA in feces by qPCR, normalized to vehicle. (**H**) Retinoic acid concentration in stool of GF and SFB mice treated with Aldh1a2 inhibitor, normalized to sample weight. Results are ± SEM. \**p* < 0.05, \*\**p* < 0.01. n.s., not significant.

### Commensal bacteria provide a direct source of retinoic acid that improves host defense

Intestinal RA levels did not reflect altered mammalian RA synthesis in SFB-colonized mice (**Figures 3E-3H**), suggesting a distinct source of RA. While microbial RA production has not been extensively studied, *Bacillus cereus* was described to express a bacterial aldehyde dehydrogenase enzyme (*bc*ALDH1A1, KFL74159.1) that produces RA *in vitro* (Hong et al., 2016). Consistent with this work, we found that *B. cereus* produced RA when incubated with the vitamin A derivative, retinol (**Figure 4A**), without altered growth (**Figure S2A**). Furthermore, retinol increased ALDH1A1 expression by *B. cereus* (**Figure 4B**), supporting that bacterial expression of this RA-producing enzyme is regulated by local retinol levels. Therefore, to investigate whether SFB and potentially other bacterial species express similar aldehyde dehydrogenases that generate RA, we compared protein sequences using *bc*ALDH1A1 as reference. Glutamate-266 and Cysteine-300 are necessary for the catalytic activity of *bc*ALDH1A1 (Hong et al., 2016).

**Figure 4:**
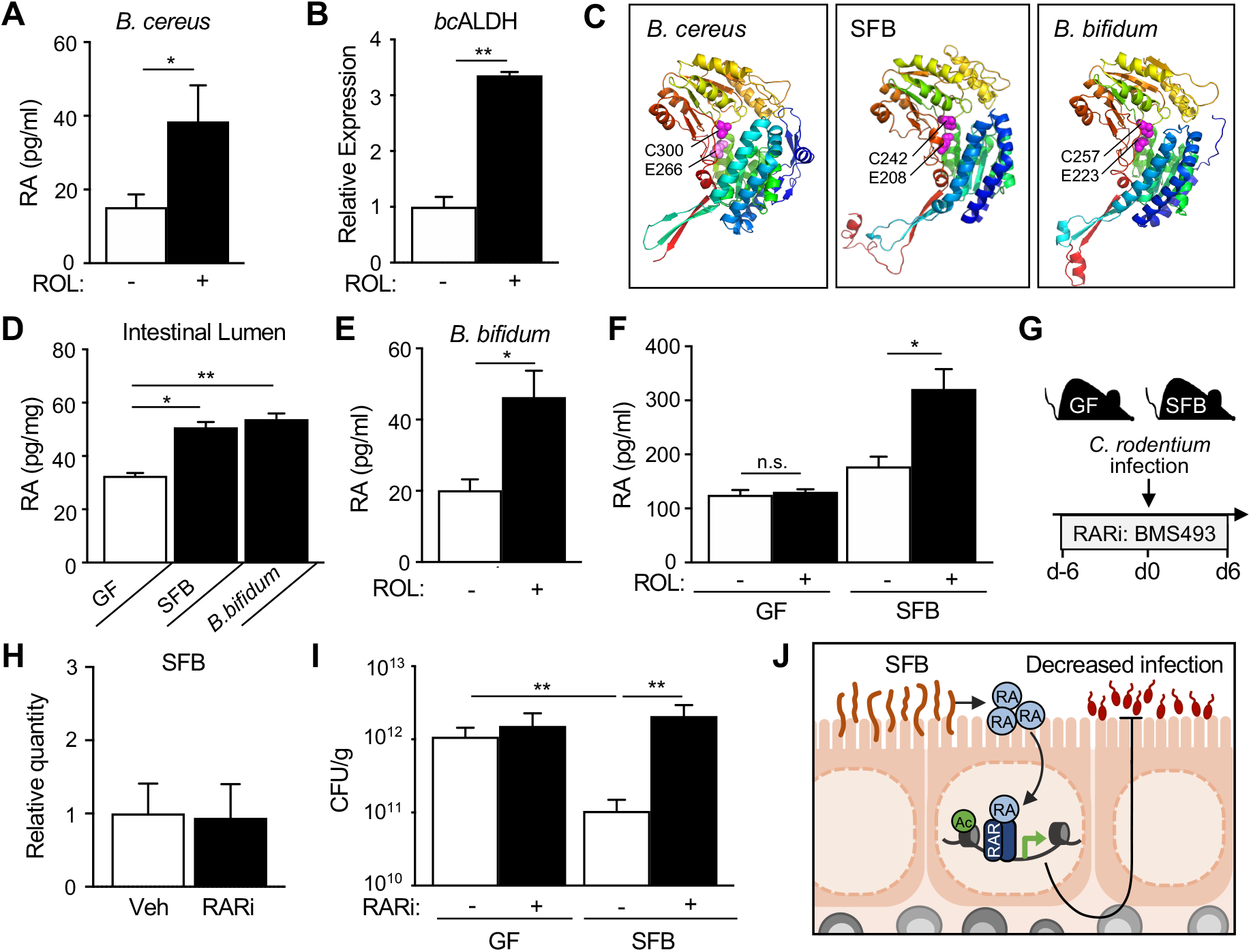
Commensal bacteria provide a direct source of retinoic acid that improves host defense. (**A**) Retinoic acid concentration in media from *B. cereus* cultured with all-trans retinol (ROL). (**B**) Expression of ALDH by *B. cereus* cultured -/+ ROL. (**C**) Protein structure of *B. cereus* ALDH1A1 (KFL74159.1), SFB ALDH (WP_007440235.1) and *B. bifidum* ALDH (WP_015438559.1). Conserved catalytic glutamate [E] and cysteine [C] residues shown as magenta-colored spheres. Root-mean-square deviation of atomic positions (RMSD) compared to *bc*ALDH1A1 for SFB ALDH=2.059 Å; *B. bifidum* ALDH, RMSD=1.514 Å. (**D**) Retinoic acid concentration in intestinal contents of GF mice monoassociated with indicated bacteria, normalized to sample weight. (**E**) Retinoic acid culture concentration following bacterial incubation with ROL, per 10^8^ CFU of bacteria. **(F**). Retinoic acid concentration in supernatants from intestine explants from GF and SFB mice incubated with vehicle or ROL. (**G**) Experimental approach. (**H**) Relative level of SFB DNA in feces by qPCR, normalized to vehicle. (**I**) *C. rodentium* CFUs in stool of GF and SFB mice -/+ RAR inhibitor (RARi; BMS493), normalized to sample weight, day 6 p.i. (**J**) Retinoic acid production by commensal bacteria represents a beneficial mechanism that enables intestinal microbiota to boost host defense against enteric infection. Results are ± SEM. \**p* < 0.05, \*\**p* < 0.01. n.s., not significant.

Interestingly, SFB, along with the well-known probiotic bacteria *Bifidobacterium bifidum*, encode bacterial ALDH enzymes that share these critical catalytic amino acid residues (**Figure 4C**). Furthermore, the predicted protein structures of ALDH enzymes from SFB and *B. bifidum* both exhibited marked overlap with *bc*ALDH1A1 (**Figure 4C**). Thus, to test whether these commensal bacterial strains generate RA *in vivo*, GF mice were colonized with SFB (**Figure S2B**) or *B. bifidum* (**Figure S2C**) and compared to GF controls. Mice colonized with ALDH-expressing SFB or *B. bifidum* demonstrated increased luminal RA relative to GF mice (**Figure 4D**). Further, culturing *B. bifidum* with retinol increased RA levels without impacting bacterial growth (**Figures 4E and S2D**). Similarly, RA was significantly increased in supernatants following retinol treatment of SFB explant cultures, whereas this induction was not observed in sterile cultures (**Figure 4F**).

Collectively, these results indicate that a subset of commensal bacterial populations, including SFB, can provide a direct source of RA in the intestine. Thus, given that SFB protects against *C. rodentium* infection (**Figure 1A**), we hypothesized that RA may mediate SFB-dependent defense against this pathogen. In order to test whether SFB promotes protection to infection by activating RAR, GF and SFB-colonized mice were treated with an RAR-inverse agonist (RARi: BMS493) (**Figure 4G**) that blocks RA-RAR activation (Metzler et al., 2018). RARi treatment did not alter SFB colonization (**Figure 4H**). Remarkably, though, impaired RA-RAR activation abrogated the protective effect of SFB against *C. rodentium*, whereas pathogen levels in GF mice were largely unaffected (**Figure 4I**). Therefore, these data demonstrate that activation of RAR signaling by commensal bacteria, such as SFB, mediates microbiota-dependent innate defense against infection (**Figure 4J**).

## Discussion

In this study, we discovered that SFB and other beneficial commensal bacteria generate RA in the intestine and regulate epithelial RAR signaling to enhance defense against a pathogen (**Figure 4J**). RA is a fat-soluble metabolite derived from carotenes and vitamin A that is known for its immunomodulatory effects and role during infectious disease (Hall et al., 2011a). Vitamin A deficiency in humans is linked to increased susceptibility to numerous bacterial and viral pathogenic infections (Sommer, 2008; World Health Organization, 2009), whereas vitamin A supplementation reduces incidence and mortality of diarrheal diseases commonly caused by intestinal pathogens (Green and Mellanby, 1928; Huang et al., 2018; Semba, 1999). Although vitamin A has been used as a clinical health intervention for diarrheal disease and other health conditions, the reported outcomes have been inconsistent (Dibley et al., 1996; Long et al., 2007). This variability has been attributed to range of factors such as varying degrees of baseline vitamin A deficiencies and differences in vitamin A dosing strategies (Dibley et al., 1996; Long et al., 2007). In mice, vitamin A availability alters severity of infection and regulates repair of infection-induced epithelial damage (McDaniel et al., 2015; Mielke et al., 2013; Snyder et al., 2019).

IECs in the small intestine express both RA-generating enzymes and RARs that are activated by RA, thereby playing a central role in vitamin A-dependent regulation. Our transcriptional analyses showed that induction of RAR targets during infection was higher with SFB-colonization and indicated RAR targets are enriched in host defense pathways. Consistent with this, recent studies have discovered that IEC-intrinsic RAR expression promotes defense by regulating antimicrobial peptide production (Gattu et al., 2019; Jijon et al., 2018). Furthermore, the ability of IECs to sense RA is necessary for defense against *Salmonella typhimurium* (Gattu et al., 2019; Iyer et al., 2020). Using a transgenic mouse model where IECs are unable to respond to RA, we observed that IEC-intrinsic activation of RAR by RA is necessary for defense against *C. rodentium.* Specifically, RA administration significantly lowered pathogen burdens in control mice compared to mice lacking the ability to activate RAR in IECs. Although we did not observe significant early protection against *C. rodentium* from RA-administration in mice with defective RAR specifically in IECs, RA also likely activates mucosal immune cells, consistent with studies showing dendritic cells, macrophages and innate-lymphoid cells also respond to RA during infection (Erkelens and Mebius, 2017; Gundra et al., 2017; Kim et al., 2015; Zeng et al., 2016).

While our studies found that SFB-dependent defense initially occurred independently from CD4^+^ T cells, both SFB and RA are known to regulate development and function of adaptive immune cells, particularly Th17 cells which promote *C. rodentium* clearance (Symonds et al., 2009). Th17 cell regulation by RA appears to be largely context and dose dependent. RA supplementation at pharmacological levels has been shown to suppress Th17 differentiation and promote regulatory T cells (Benson et al., 2007; Mucida et al., 2007). In contrast, others have found that physiological concentrations of RA stimulation instead promote Th17 skewing of naïve CD4^+^ T cells both *in vitro* and *in vivo* (Takahashi et al., 2012; Uematsu et al., 2008; Wang et al., 2010). Consistent with this evidence, RARα-deficient T cells are also unable to differentiate into Th17 cells *in vitro* under Th17-polarizing conditions (Hall et al., 2011b). Importantly, mice fed vitamin-A deficient diet are deficient in Th17 cells, further indicating that RA is required for *in vivo* development and/or maintenance of these cells (Cha et al., 2010; Wang et al., 2010). The presence of SFB in the intestinal microbiota of mice was shown to drive Th17 cell expansion by inducing epithelial expression of Serum Amyloid A (SAA) proteins (Ivanov et al., 2009; Sano et al., 2015). Intestinal SAA expression requires dietary vitamin A and is directly regulated by epithelial RARs (Gattu et al., 2019).

These findings, in combination with our current work, support a model in which increased RA levels and enhanced epithelial RAR activation by SFB promotes innate epithelial defense, and simultaneously drives Th17 cell differentiation, potentially through RA-dependent transcriptional regulation of SAAs. Considering that SFB directly interacts with IECs and even undergoes vesicle-mediated communication (Ladinsky et al., 2019), we suspect that SFB may provide a local dose of RA that transcriptionally primes adherent cells. SFB colonizes the terminal ileum, however *C. rodentium* infects the colon. This spatial separation suggests that RA from SFB may either regulate IECs located in the more distal intestine and/or indirectly enhance innate defense in the colon.

Despite the abundance of evidence linking host immunity and RA, relatively little is known about how the microbiota regulates RA. It was recently shown that intestinal tissue RA levels were lower in conventionally-housed mice compared to GF mice due to decreased expression of Rdh7, an enzyme that oxidizes retinol to retinal (Grizotte-Lake et al., 2018). This downregulation was found to be driven primarily by Clostridial species. Furthermore, expansion of proteobacteria following antibiotic depletion of Clostridium correlated with induction of host Rdh7 expression, suggesting that different commensal bacterial species may differentially regulate RA. Our data describe a distinct mechanism with SFB. Intestinal RA levels increased in SFB-colonized mice compared to GF, however host RA-producing gene expression remained similar in IECs from SFB colonized mice. Given this regulation, we anticipate that commensal bacterial species likely differentially modulate intestinal RA levels through microbe-intrinsic mechanisms and/or distinct host-dependent pathways.

Investigation of vitamin A metabolism has largely focused on mammalian enzymes, as the RA production has generally been considered a mammal-specific reaction (Biesalski et al., 2007). Whether bacteria directly contribute to vitamin A metabolism has been largely unexplored. Prior studies demonstrated that *E. coli* were capable of generating retinal and retinyl acetate, and potentially retinoic acid, in culture (Jang et al., 2011, 2015). Genetic manipulation of putative endogenous genes in *E.coli* involved in converting retinol to retinal *(ybbo)* and RA *(puuC, eutC)* altered retinoid production by *E. coli* (Jang et al., 2011, 2015). Furthermore, a bacterial ALDH expressed in *B. cereus*, a gram-positive bacterium commonly found in the gastrointestinal tract of mammals, was able to directly convert retinal to RA *in vitro* (Hong et al., 2016). Together these findings indicate that bacteria inherently harbor retinoid metabolism pathways.

In addition to the protective effects of bacterial RA on the host, we anticipate that there are likely bacterial-intrinsic benefits to metabolizing vitamin A. While we did not observe obvious differences in cultured bacterial growth with short-term vitamin A exposure, it is possible that metabolizing vitamin A entails a competitive advantage in the intestine. Vitamin A availability affects cellular zinc absorption, and vice versa (Christian and West, 1998; Rahman et al., 2002; Smith, 1980). Zinc is an essential micronutrient for all organisms including bacteria and is required for normal cellular physiology. However, excess zinc is toxic to bacteria and thus must be tightly controlled (Hantke, 2005; McDevitt et al., 2011). Zinc is most abundant in the intestine, so it is plausible that bacterial metabolism of retinoids improves absorption of zinc to maintain non-bactericidal levels in the intestinal environment. Alternatively, oxidation of vitamin A to retinol may provide bacteria with important reducing equivalents in the form of NADH and NADPH that are needed for energy metabolism (Spaans et al., 2015; Sporer et al., 2017).

Collectively, this work reveals a new level of regulation in which microbiota-derived RA enhances host defense to a bacterial pathogen. To our knowledge, the ability of commensal bacteria, such as SFB, to provide a direct source of RA in the intestine that physiologically regulates the mammalian host has not been described. Dissecting upstream and downstream factors in this new pathway will form the basis for further investigations into the mechanistic underpinnings for commensal bacterial RA production. In addition to *B. cereus* and SFB, we found that microbial RA regulation extended to another beneficial bacterial strain, the probiotic *Bifidobacterium bifidum*. Thus, these findings support development of pro- and prebiotic approaches that induce elevated intestinal RA in order to decrease enteric infection.

## Acknowledgements

We thank A. Herr, and the Way, Qualls, Haslam, and Deshmukh labs for useful discussions and members of the Alenghat lab for critical reading of the manuscript. We thank CCHMC Veterinary Services, Research Flow Cytometry Core, Pathology Research Core, and S. Jagannathan, and A. Barski for services and technical assistance. We thank the Yakult Central Institute for providing SFB. This research is supported by the National Institutes of Health (DK114123, DK116868 to T.A., and F32AI147591 to E.M.E.), Cardell Fellowship to V.W., Pew Charitable Trust, and a Kenneth Rainin Foundation award to T.A. T.A. holds an Investigator in the Pathogenesis of Infectious Disease Award from the Burroughs Wellcome Fund. This project is supported in part by PHS grant P30 DK078392 and the CCHMC Trustee Award and Procter Scholar’s Program.

## Author Contributions

Conceptualization - V.W., T.A.; Methodology - V.W., E.M.E., B.V., T.A.; Investigation - V.W., E.M.E., J.W., S.H.H., S.W., L.E., T.R.; Formal Analysis - V.W., R.K., E.M.E.; Writing – Original Draft - V.W., T.A.; Writing, Review & Editing - V.W., E.M.E., S.H.H., T.A.; Funding Acquisition - T.A., E.M.E., V.W.; Resources - B.V.; Supervision – T.A.

## Methods

### Mice

Germ-free (GF) mice were maintained in sterile isolators (Class Biologically Clean) or sealed positive pressure IVC racks (Allentown) in the CCHMC Gnotobiotic Mouse Facility. For monoassociation studies, GF mice were colonized with singular commensal species suspended in sterile PBS via oral gavage (*Bacillus cereus* ATCC 10786; *Bifidobacterium bifidum* ATCC 29521) or by pre-colonized bedding (SFB). All germ-free and monoassociated mice were fed autoclaved food and water, and routinely monitored to ensure the absence of microbial contamination and/or assess level of colonization. C57BL/6 floxed dnRAR (Rajaii et al., 2008) mice were crossed to villin-Cre-recombinase expressing mice (Madison et al., 2002) to generate dnRAR^IEC^ mice. Animals were housed in ventilated cages up to 4 per cage in 12 hr light/dark cycles with unrestricted access to food and water. All mouse experiments were conducted according to the Institutional Animal Care and Use Committee (IACUC). Animals were cared for by a licensed veterinarian and proper steps were taken to ensure the welfare and minimize the suffering of all animals in the conducted studies.

### *C. rodentium* infections

Mice were orally infected with 10^9^ colony-forming units (CFUs) of *C. rodentium* suspended in sterile PBS. Post-infection CFUs were measured in stool homogenized in 500ul PBS in a Tissue Lyser II at 30 Hz for 3 min. Homogenates were serially diluted 10-fold on MacConkey agar and CFUs were counted after 16 hr incubation at 37’C, normalized to fecal weight. For RA studies, mice were orally gavaged with 300μg all-trans retinoic acid (Sigma) or vehicle (DMSO) in 100μl corn oil q.d. 5 days prior to and during the infection. For RAR inhibitor (RARi) studies, 400μg BMS493 (Torcis Bioscience) suspended in 10% DMSO/corn oil was administered to mice via oral gavage q.o.d. over 6 days pre-infection and 6 days post-infection. For Aldh1a2 inhibition, mice were orally gavaged with 125 mg/kg of WIN18446 (R&D Systems) or vehicle (DMSO) in 100ul corn oil for 8 days q.o.d.

### CD4^+^ T cell depletion and flow cytometry

CD4^+^ cells were depleted using anti-CD4 monoclonal depletion antibody (clone: GK1.5) or matching isotype control (Rat IgG2B) administered intraperitoneally, 500μg per day every 3 days for a total of 3 doses. Efficacy of CD4^+^ depletion was determined in colonic lamina propria and spleen by flow cytometry. For intestinal lamina propria lymphocytes isolation, tissue pieces were washed with cold PBS and incubated in RPMI with 1 mg/ml Collagenase/Dispase for 30 min at 37’C with shaking at 200 rpm. Splenocytes were disrupted into single cell suspension by passing the organ through 70 μm filter and RBCs were lysed in ACK lysis buffer (Invitrogen) for 3 min.

Cells were stained using the following monoclonal fluorescence-conjugated antibodies: BUV395 anti-CD45.2 (Clone: 104, BD Biosciences), APC-eFluor 780 anti-CD4 (Clone: RM4-5, eBioscience), and APC anti-CD8a (Clone: 53-6.7, eBioscience). All antibodies were diluted in FACS buffer (2% FBS, 0.01 Sodium Azide, PBS). Dead cells were gated out by using the Fixable Violet Dead Cell Stain Kit (Invitrogen). Samples were acquired on the BD LSRFortessa (BD Biosciences) and analyzed with FlowJo Software (Treestar).

### IEC isolation and RNA analyses

IECs were isolated from distal small intestine by shaking 10 cm of tissue in 1mM EDTA/1mM DTT 5% FBS at 37’C for 10 min as described previously (Alenghat et al., 2013). Bacteria were treated with RNAprotect Bacteria Reagent (Qiagen) for 5 min prior to RNA isolation. RNA was extracted from cells using the RNeasy Kit (Qiagen) according to manufacturer’s instructions. For RT-qPCR, RNA was reverse-transcribed with Verso reverse transcriptase (Invitrogen) and expression was compared using SYBR (Applied Biosystems) and analyzed in the linear range of amplification. Target gene expression was normalized to an unaffected control gene. For global expression analyses, 3-4 biological replicates of IECs from *C. rodentium-infected* GF and SFB-monoassociated mice were compared. Following removal of primers and barcodes, raw reads were processed using Kallisto, which employs pseudoalignment to assess compatibility between raw reads and genomic targets. Annotations were provided by UCSC with transcripts per million (TPM) as output, which were log2-transformed and baselined to the median of all samples. Further, transcripts were filtered to include only those with TPM >3 in 100% of samples in at least one condition. Differential expression was assessed with a moderated t-test with *p*<0.05 and fold-change>1.5. For gene ontology analyses, differential gene lists were submitted to DAVID bioinformatics database (david.ncifcrf.gov). Pathway enrichment significance are displayed as log10-transformed *p*-values.

### ChIP-seq

ChIP-seq on IECs was performed as described previously (Wu et al., 2020) with a few modifications. Briefly, cells were fixed for 10 min in 1% PFA at room temperature, followed by quenching with 125mM glycine for 10 min. After a two-step wash with cold PBS, fixed cells were lysed, and nuclear extracts were washed in TE 0.1% SDS with protease inhibitors and sonicated using a S220 Focused-ultrasonicator (Covaris). Prior to immunoprecipitation, sheared chromatin was precleared for 20 min at 4’C using Protein G Dynabeads (Thermo Fisher Scientific). Immunoprecipitations were performed using fresh beads and anti-Histone H3 acetyl K27 (H3K27Ac) antibody (Abcam: ab4729) using a SX-8G IP-STAR automated system (Diagenode) with the following wash buffers: (1) RIPA 150mM NaCl, (2) RIPA 250mM NaCl, (3) LiCl 250mM, 0.5% sodium deoxycholate, NP40 0.5%, and (4) TE 0.2% Triton X-100. Immunoprecipitated chromatin were treated with Proteinase K (Thermo Fisher Scientific) at 42’C for 30 min, 65’C for 4 hr, and 15’C for 10 min in elution buffer (TE 250mM NaCl 0.3% SDS). Phenol:chloroform isoamyl alcohol with Tris-HCl (pH 8.0) and chloroform phase-separation were used to isolate DNA, followed by overnight ethanol precipitation. ChIP DNA was sequenced using Illumina HiSeq 2500 platform. ChIPseq data were processed using analytic pipelines in galaxy (usegalaxy.org). Following raw read alignment to mm10, MACS2 was used for peak calling and differential peak detection. Peaks were visualized by the UCSC genome browser in Biowardrobe (Kartashov and Barski, 2015). Transcription factor-binding site motifs were identified within 150 bp of the center of the differential peaks using PscanChIP (JASPAR 2018 database), displayed as the global *p*-value.

### Immunofluorescent tissue staining

Distal large intestine was fixed in 4% paraformaldehyde overnight at 4’C and then placed in 30% sucrose for 24 hr. Tissues were embedded in OCT compound and cut as frozen sections (10 μm). Frozen sections were thawed and blocked with 1% BSA for 1 hr at room temperature. The following antibodies were diluted in 0.5% BSA and incubated with the tissue for 1.5 hr at room temperature: Alexa Fluor 488-anti-GFP (5.0 ug/ml, A21311, Invitrogen) and Alexa Fluor 594-Phalloidin (1:200, A12381, Invitrogen). Nuclei were stained with DAPI (4’,6-Diamidino-2-Phenylindole, Dihydrochloride, 0.5 ug/ml). Slides were washed and then mounted using Fluoromount-G (Invitrogen 00-4958) and imaged on a Nikon A1R LUN-V inverted confocal microscope.

### Bacterial culture and quantification

*C. rodentium* were cultured *in vitro* in 96-well round bottom plates with DMSO, 1nM-10μM RA or 40 μg/ml Kanamycin (Sigma) at 37’C shaking at medium speed in a microplate reader (Biotek Synergy 2). Bacterial density (OD600) was measured every hour over 16 hr. For bacterial retinol culture studies, bacteria were grown in liquid cultures overnight at 30’C or 37’C at 180 rpm for 16 hr. Bacterial suspensions were then diluted 1:3 in PBS and incubated with 1μM all-trans retinol (Sigma) for 3 hr in a 24-well plate at 37’C with gentle shaking at 120 rpm under light-restricted conditions. To determine bacterial levels, fecal or cultured bacterial DNA was isolated using QIAamp Fast DNA Stool Mini Kit (Qiagen) following the kit protocol. Bacterial DNA was assessed by quantitative PCR (QuantStudio3; Applied Biosystems) using bacterial-specific or 16s primer pairs (Invitrogen, MilliporeSigma).

### Retinoic acid quantification

Intestinal contents and IECs pellets were collected under dark conditions and homogenized in PBS. Extracts were run on a retinoic acid ELISA kit (MyBiosource, MBS706971) according to manufacturer instructions. Briefly, samples were incubated with 50 ul HRP-conjugated antibody for 40 min at 37’C, washed 5 times with wash buffer, and incubated with TMB substrate for 20 min at 37’C. The reaction was quenched, and absorbance was measured of each well using a microplate reader (Biotek Synergy 2) set to 450 nm. For explant experiments, equal sections of terminal ileum were taken from GF and SFB-monoassociated mice and cultured in a 24-well plate with 1μM all-trans retinol (Sigma) for 3 hr at 37’C without light. RA was measured in culture supernatant after incubation.

### Protein modeling and alignment

To predict 3D structures, protein sequences were submitted to the Phyre2 server (http://www.sbg.bio.ic.ac.uk/phyre2) (Kelley et al., 2015) and modeled after existing Protein Data Bank templates (bcALDH1A1: PDB c4pt3C; SFB ALDH: PDB c6k0zA; *B. bifidum* ALDH: PDB c4f9iA). Figures were generated using the PyMOL Molecular Graphics System, Version 2.4 Schrodinger, LLC (https://pymol.org/). The superimpose function was used to determine structural similarity to *bc*ALDH1A1, reported as the overall root-mean-square deviation (RMSD) value.

### Statistics

All statistical analyses were performed using GraphPad Prism 8.0. Statistical significance was determined by students t-test or ANOVA. All data meet the assumptions of the statistical tests used. Results are shown as mean ± SEM and considered significant at *p*<0.05 (*); *p*<0.01 (**).

