## Supplemental Figures for "Commensal bacterial-derived retinoic acid primes host defense to intestinal infection"

### Supplemental Figure S1

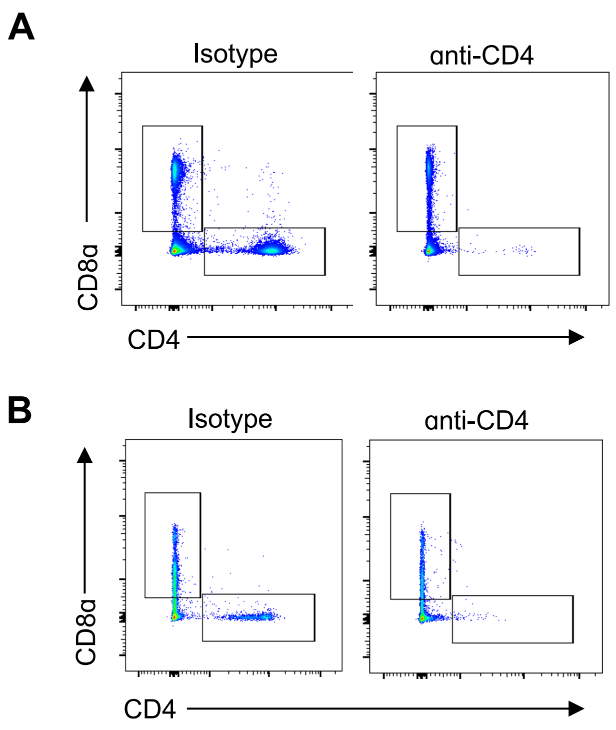


**Figure S1: CD4⁺ T cell depletion. (A, B**) Representative flow cytometry plots for (A) spleen and (B) colon from isotype and anti-CD4 depleted mice. Gated on CD45⁺ cells.

Supplemental Figure S2

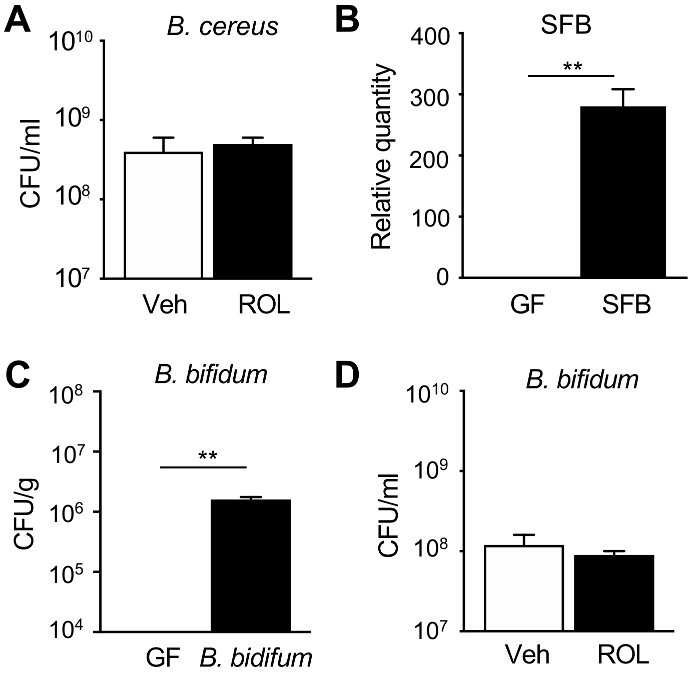


**Figure S2: Inhibition of host RA production or signaling does not affect intestinal SFB levels.** (**A**) CFUs of *B. cereus* grown *in vitro* -/+ retinol (ROL). (**B**) SFB DNA in feces by qPCR. (**C, D**) CFUs of *B. bifidum* in (C) feces from *B. bifidum*-monoassociated GF mice or (D) grown *in vitro.* Results are ± SEM. ***p*< 0.01.
